# Comprehensive characterization of skeletal muscle remodeling in *hSOD1^G93A^* mice reveals limited functional impact of systemic FOXO1 inhibition

**DOI:** 10.64898/2026.05.27.726208

**Authors:** Ainhoa Vidal-Gil, Ignacio Azcue, Maria Levchuk, Amaia Elicegui, Oihane Pikatza-Menoio, Maddalen Robles-Cantero, Ane Otegui, María Rodríguez-Hidalgo, Laura Moreno-Martínez, Cristina Ruiz-Roldan, Andrea Valls, Bahaa Daou, Mikel García-Puga, Itziar Vergara, Ander Matheu, Amets Saenz, Rosario Osta, Adolfo López de Munain, Sonia Alonso-Martín

## Abstract

**Background:** Amyotrophic lateral sclerosis (ALS) is a fatal neurodegenerative disorder characterized by progressive motor neuron (MN) loss, muscle atrophy and paralysis. Although traditionally considered a MN-specific disease, accumulating evidence supports a crucial contribution of skeletal muscle pathology to disease onset and progression. Except for specific mutations, to date there is no effective treatment for ALS. FOXO transcription factors regulate programs of atrophy, metabolism and stress response in skeletal muscle, and their inhibition has shown beneficial effects in cellular and *Drosophila* models of ALS.

**Methods:** In this study, we investigated whether pharmacological FOXO inhibition (iFOXO) could modify disease progression and muscle pathology in female *hSOD1^G93A^* mice. Mice received daily oral administration of iFOXO starting at presymptomatic (P50; n=5 per group) or symptomatic (P90; n=9 mice per group) stages until end-stage. Body weight was monitored longitudinally, and motor performance was evaluated using grip strength and hanging-wire tests. *Tibialis anterior* and *soleus* muscles, representing fast- and slow-twitch muscles respectively, were analyzed by histology and immunofluorescence to assess fiber atrophy, fibrosis, lipid accumulation, satellite cell pool and fiber type composition. Quadriceps muscles (n=3 per group) were used for RNA-seq analysis.

**Results:** While histological analyses revealed severe fiber atrophy and increased fibrosis in *hSOD1^G93A^* mice, satellite cell numbers were preserved or mildly increased in a muscle and treatment onset dependent manner. iFOXO treatment did not improve motor performance, survival or attenuate muscle atrophy. Transcriptomic profiling indicated that genotype was the predominant driver of gene expression changes, while iFOXO produced only subtle, treatment onset dependent effects on pathways related to oxidative stress responses, mitochondrial function and adaptive metabolism.

**Conclusion:** Overall, FOXO inhibition alone showed limited therapeutic benefit in the *hSOD1^G93A^* ALS mouse model. These findings highlight the dominant influence of ALS driven molecular alterations over pharmacological modulation and emphasize the need for combinatorial therapeutic strategies targeting multiple disease mechanisms, including those preserving nerve health.

## INTRODUCTION

Amyotrophic lateral sclerosis (ALS) is a devastating neurodegenerative disease characterized by progressive loss of upper and lower motor neurons (MN), leading to a rapid decline of motor function, muscle paralysis and death due to respiratory failure, typically within 3-5 years of symptom onset. The etiology of the disease remains unknown, and no effective treatment is currently available (1). With approximately 90% of sporadic cases, more than 40 genes have been associated with ALS (2), with the most frequent mutations occurring in *C9orf72*, *TARDBP* (encoding TDP-43), *SOD1* and *FUS* (3,4).

ALS has been considered as a MN-specific disease, however, it is now recognized as a multisystem disorder involving structural, physiological, and metabolic alterations in several cell types that contribute to disease onset and progression (5,6). Increasing evidence from clinical and experimental studies indicates that MN degeneration may arise from non-cell-autonomous mechanisms involving glial (5,7) and muscle cells (5,8,9). Notably, overexpression of mutant SOD1 exclusively in MNs does not recapitulate ALS pathology, whereas its modulation in skeletal muscle is sufficient to induce motor deficits, muscle atrophy and MN degeneration (10,11). These findings support a *dying back* model in which pathological alterations originate distally in skeletal muscle and propagate retrogradely through the neuromuscular junction (NMJ) before MN degeneration and symptoms onset (5,12). Consistently, myotubes derived from ALS patients secrete neurotoxic vesicles that induce MN death *in vitro* (13) and *FUS* mutant MN and myotubes co-culture disrupts NMJ formation (14,15). Together, these findings highlight the contribution of muscle-derived pathological signals to ALS progression.

Skeletal muscle possesses a remarkable regenerative capacity, even after severe injury (16). However, accumulating evidence suggests that this capacity is impaired in ALS. Myoblasts isolated from ALS patients or animal models display reduced differentiation potential and altered myogenic gene expression (17–19). Primary myoblasts from ALS patients also exhibit metabolic dysfunction and increased sensitivity to oxidative stress (20), indicating that intrinsic muscle abnormalities may contribute to disease progression. Although the underlying causes of ALS are heterogeneous, metabolic dysfunction is a shared hallmark, with energy homeostasis disruption negatively affecting disease severity and survival (3). Strikingly, transcriptomic analyses of patient derived myoblasts have revealed enrichment of pathways associated with glucose metabolism and insulin signaling, along with increased transcriptional binding sites for FOXO transcription factors (19). These findings suggest that skeletal muscle is not only an active contributor to ALS pathology but also represents a promising therapeutic target.

Currently, ALS treatments are limited and mainly aim to slow disease progression or alleviate the symptoms. Disease modifying therapies include Riluzole or Edaravone, modestly extending survival or slowing the rate of functional decline, respectively (21,22). More recently, Tofersen, an antisense oligonucleotide targeting SOD1, modifies disease progression by reducing SOD1 protein levels, with some of the patients demonstrating improvements in muscle strength (23). However, this treatment is restricted to patients carrying SOD1 mutations. Overall, these therapies offer limited benefits and highlight the urgent need for more effective treatment strategies.

FOXO transcription factors (FOXO1, FOXO3 and FOXO4) are key regulators of skeletal muscle homeostasis, coordinating processes such as protein turnover, autophagy, stress responses and metabolism. Under catabolic or pathological conditions, FOXO activation induces the expression of ubiquitin ligases such as *Fbxo32* (atrogin-1) and *Trim63* (MuRF1), promoting proteolysis and muscle atrophy (24,25). FOXO3 also regulates autophagy related genes such as *Gabarapl1* and *Ctsl*, thereby promoting autophagy during muscle wasting (26,27). Beyond proteostasis, FOXOs modulate energetic metabolism, oxidative stress defense and fiber type specification (28,29). Importantly, pharmacological or genetic inhibition of FOXO signaling has been shown to preserve muscle mass and function in models of disuse and cachexia (30,31), highlighting FOXO as a potential therapeutic target in conditions involving muscle degeneration and metabolic stress.

We have previously identified FOXO1 as a key mediator of intrinsic metabolic and myogenic defects in ALS patients’ myoblasts. Furthermore, we demonstrated that pharmacological inhibition of FOXO1 (iFOXO), through inhibition of FOXO1 transcriptional activity and downstream gene regulation, improved muscle differentiation capacity and improved metabolic parameters *in vitro*, while ameliorating motor deficits and extending survival in *Drosophila melanogaster* muscle-specific TDP-43 and FUS loss of function models (19).

Aiming at reversing the muscle pathophysiology observed in the *hSOD1^G93A^*mouse model, we conducted iFOXO treatments. We have characterized histopathological hallmarks of skeletal muscle in *hSOD1^G93A^* mice following iFOXO treatment either at presymptomatic and symptomatic stages of disease progression. This dual intervention enables to evaluate both the preventive and the therapeutic potential of FOXO inhibition in ALS. Given the distinct metabolic profiles of muscle fiber types, with fast-twitch fibers being more glycolytic and vulnerable in ALS, whereas slow-twitch fibers are more oxidative and relatively preserved, we analyzed *tibialis anterior* (TA) and *soleus* muscles to capture fiber-type specific alterations (32). In summary, deep analysis of the skeletal muscle demonstrated an atrophic phenotype towards smaller myofibers and increased fibrosis in both fast (TA) and slow (*soleus*) muscles in terminal animals, with a clear switch towards slow myofibers, with a reduction on the proportion of fast fibers in both muscles. Strikingly, there is a trend of increased numbers of the muscle stem cell population in the *soleus*, which is further enhanced when treatment initiated at the presymptomatic stage. Pathway analyses reveal that SOD1 mutant muscle is characterized by impaired energy metabolism and contractile function together with activation of stress and myogenic programs, reflecting a state of chronic muscle stress. iFOXO treatment modulates these responses in a treatment onset-dependent manner but fails to restore muscle structure or function. Together, though implying multitarget treatments for heterogenous and complex diseases such as ALS, we present an exhaustive characterization of the skeletal muscle tissue in the *hSOD1^G93A^*mouse model at end-stage of the disease.

## MATERIALS AND METHODS

### Animals and ethics

The transgenic *hSOD1^G93A^* mouse strain (*B6SJL-Tg(SOD1*G93A])1Gur/J*; Jackson Laboratoy, Strain #:002726) was used as ALS disease model. Age-matched non-transgenic wild-type (WT) littermates were used as controls. Genotyping was performed by polymerase chain reaction using DNA extracted from tail biopsies to identify hemizygous transgenic and WT animals. Mice were maintained under standard conditions, a 12/12 h light/dark cycle and *ad libitum* access to food and water. All experimental procedures were approved by the Biogipuzkoa Health Research Institute Ethics Committee in agreement with Spanish and European regulation on animal welfare: PRO-AE-SS-193 (OH-20-37) and PRO-AE-SS-238 (OH-21-42), in line with the NC3R (National Centre for the Replacement, Refinement and Reduction of Animals in Research) guidelines. In survival studies, the humane endpoint was defined as the loss of the righting reflex for more than 30 seconds (s). Afterwards, animals were anesthetized and sacrificed using CO_2_ and muscles from both hindlimbs and forelimbs were collected.

### Therapeutic efficacy of iFOXO administration in *hSOD1^G93A^* mice

A dose of 20 mg/kg of FOXO1 inhibitor AS1842856 (S8222, Astella Pharma, Shelleck) was administered by oral *gavage* once daily until end-stage of the disease (∼4-month-old). *hSOD1^G93A^* female mice were randomly distributed to treatment groups matched by age and litter. Two treatment regimens were established: one starting at postnatal day 90 (P90; n=27, 9 mice per treatment group) and another at P50 (n=15, 5 mice per treatment group). Control mice received equivalent volumes of vehicle (2-Hydroxypropyl)-β-cyclodextrin (HP-b-CD; H107, Sigma-Aldrich).

### Behavioral assays

Behavioral assessments were carried out to evaluate the effects of iFOXO treatment on locomotor function and survival in *hSOD1^G93A^*mice. All analyses were performed under blinded conditions with respect to treatment allocation. Body weight was monitored longitudinally alongside behavioral analysis.

#### Hanging-wire test

Muscular strength was evaluated using the hanging-wire test. The device was custom-built from a commercially available rectangular plastic lid (30×40 cm). The central section of the lid was removed to create an open rectangular frame. A metal wire mesh (1 mm wire diameter, 1×1 cm grid spacing) was then securely attached to the opening. To perform the hanging-wire test, mice were placed on an inverted wire grid and the latency to fall was recorded. Each mouse performed two trials, with a maximum duration of 120 s, and the longest latency was used for analysis.

#### Grip strength test

Forelimb and hindlimb grip strength was assessed using a grip strength meter (BIOSEB, EB Instruments). Mice were habituated to the procedure one week prior to treatment initiation. During testing, animals grasped the metal grid with all four paws while being gently pulled by the tail until release. Five trials were performed per mouse, and the mean of the three highest values was used for statistical analysis. Data were normalized with mice body weight.

### Histological analysis

The TA and *soleus* muscles were snap-frozen in liquid nitrogen-cooled isopentane for subsequent histological analysis (19). Cryosections of 8 µm thickness were obtained and processed following previously described protocols for Hematoxylin and Eosin (19,33), Oil Red O (34) and Picrosirius Red staining (34). Histological sections were mounted using DPX Mountant (06522, Sigma-Aldrich) and scanned at 20X magnification using the ZEISS AxioScan 7 slide scanner.

### Immunofluorescence

Cryosections (8 µm) of TA and *soleus* muscles were processed differently depending on the target marker. For satellite cell staining, sections were fixed in 4% PFA (15713S, Electron Microscopy Sciences) for 10 minutes (min), followed by PBS washes and permeabilization for 12 min in 0.5% Triton X-100 (0694, Amresco) in PBS. Blocking was then performed for 3 hours (h) at room temperature (RT) in a solution containing 4% BSA (001-000-162, Jackson ImmunoResearch), 1% normal goat serum (NGS, 005-000-121, Jackson ImmunoResearch), 0.025% Tween-20 (822184, Sigma-Aldrich) and 0.1% sodium azide. Sections were incubated overnight at 4 °C with primary antibodies diluted in blocking solution: mouse anti-Pax7 (sc-81678, 1:50, Santa Cruz) and rabbit anti-Laminin (L9393, 1:300, Sigma-Aldrich). For muscle fiber type staining, sections were not fixed. Permeabilization was performed during the blocking step for 1 h at RT using 4% BSA, 1% NGS, 0.5% Triton X-100 and 0.1% sodium azide. Primary antibodies against myosin heavy chain (MyHC) isoforms were applied sequentially: mouse IgG2b anti-MyHC I (BAD5, 2:3, DSHB) and mouse IgG1 anti-MyHC IIa (SC-71, 1:3, DSHB) overnight at 4 °C and mouse IgM anti-MyHC IIb (BF-F3, 2:3, DSHB) and rabbit anti-Laminin (L9393, 1:300, Sigma-Aldrich) for 2 h at RT the following day.

After incubation with primary antibodies, all sections were washed in PBS and incubated for 1 h at RT in the dark with appropriate Alexa Fluor Plus or DyLight-conjugated secondary antibodies: goat anti-mouse IgG1 488 (A21121, 1:500, Invitrogen), goat anti-mouse IgG1 647 (A21240, 1:300, Invitrogen), goat anti-mouse IgG2b 555 (A21147, 1:300, Invitrogen), goat anti-mouse IgM 488 (A21042, 1:300, Invitrogen), goat anti-rabbit 647 (A21246, 1:500, Invitrogen) and goat anti-rabbit 405 (111-475-003, 1:100, Jackson ImmunoResearch). 4′,6-diamidino-2-phenylindole (DAPI) was included for nuclear counterstaining. Slides were mounted with aqueous mounting medium and imaged using ZEISS AxioScan 7 slide scanner and Axio Observer 7 systems.

### RNA-seq analysis

#### RNA preparation and RNA-seq performance

Total RNA was extracted from the quadriceps muscle (n=3 per group) using the Maxwell RSC simplyRNA Tissue Kit (AS1340, Promega). RNA concentration was determined using fluorometric quantification on a Qubit 4 instrument (Thermo Fisher Scientific), and RNA integrity was evaluated with the Agilent RNA 6000 Nano Kit on a 2100 Bioanalyzer (Agilent Technologies). All samples displayed high RNA quality, with RNA integrity numbers (RIN) greater than 7. mRNA molecules were selectively isolated and purified using the Dynabeads™ mRNA Purification Kit (Thermo Fisher Scientific). Strand-specific RNA libraries were constructed using the MGIEasy Fast RNA Library Prep Set (MGI) on the MGISP100 platform, and library quality was assessed with the Agilent High Sensitivity DNA Kit on the 2100 Bioanalyzer. Next-generation sequencing was performed on a DNBSEQ-G400 platform, generating paired-end 150 bp reads with roughly 40 million reads per sample. All procedures were carried out by Dreamgenics.

#### RNA-seq data processing and analysis

RNA-seq data processing and differential expression analysis were performed on the Genomic Platform of the Biogipuzkoa HRI using the standardized nf-core pipelines nf-core/rnaseq (v3.15.0) and nf-core/differentialabundance (v1.5.0) (35), ensuring reproducibility and adherence to best practices in computational workflows. Raw sequencing data were processed using nf-core/rnaseq. Low-quality bases filtering (Phred score < 15) using a sliding window approach and adapter were performed using fastp v.0.23.4 (36). Ribosomal RNA (rRNA) was removed with SortMeRNA v.4.3.6 (37). The resulting clean reads were then aligned to the *Mus musculus* GRCm39 reference genome (ENSEMBL) using the STAR aligner v.2.7.10a (38). The full sequence for the *Homo sapiens SOD1* gene (ENSEMBL ID: ENSG00000142168) was concatenated to the mouse reference genome index prior to alignment to enable accurate quantification of the human transgene. Transcript abundance was quantified using Salmon v.1.10.1 and subsequently aggregated to the gene level (39). Count matrices generated by nf-core/rnaseq were subjected to differential expression analysis using nf-core/differentialabundance, which employs DESeq2 (v1.44.0) (40). *p*-values were adjusted with the Benjamini–Hochberg (BH) method to reduce the number of false positives. Differentially expressed genes were called with thresholds of |log2(fold change)| ≥ 1 and adjusted *p*-value (FDR) < 0.05. Over-representation analysis (ORA) was performed using the enrichGO function from the R package clusterProfiler (v.4.12.6) (41). Differentially expressed genes were separated into upregulated and downregulated gene sets, and Gene Ontology (GO) enrichment analysis was performed independently for each subset. Only significant enrichment results (adjusted *p*-value ≤ 0.05) were considered. GO terms of interest were selected from the enrichment results and the top 15 terms were visualized.

### Statistical analysis

Data are presented as mean ± standard deviation (SD). Normality of data distribution was assessed using the Shapiro-Wilk test (*p*<0.05). Homogeneity of variances was evaluated using Bartlett’s test. For normally distributed data with equal variances, one-way ANOVA followed by Tukey’s *post hoc* test was applied. When variances were unequal, Welch ANOVA followed by Games–Howell *post hoc* test was used. For non-normally distributed data, the Kruskal-Wallis test followed by Dunn’s *post hoc* test was applied. Survival curves were generated using the Kaplan–Meier method, and differences between groups were assessed using the log-rank (Mantel-Cox) test. Statistical significance was defined as *\*p*<0.05*, **p*<0.01, and *\*\*\*p*<0.001.

## RESULTS

### Effects of iFOXO on locomotor performance and survival in *hSOD1^G93A^* mice

Our previous *in vitro* data demonstrated that inhibition of FOXO transcription factors improved ALS patient-derived myoblast differentiation as well as metabolic fitness (19). We used the quinolone derivate AS1842856 (iFOXO) in a *Drosophila* ALS model as well, improving their locomotor and survival rates (19). Thus, we next decided to treat *in vivo* the *hSOD1^G93A^*, the most used ALS animal model, initiating treatment at P90, a symptomatic stage that reflects a typical clinical scenario at diagnosis and at P50, a presymptomatic stage to assess potential preventive effects (Figure 1a). For this clinical relevance, analyses are first presented for the symptomatic intervention at P90, followed by the presymptomatic P50 treatment. Daily administration of iFOXO by oral *gavage* until end-stage (higher in females, >P150) did not improve the characteristic motor deficits of *hSOD1^G93A^*mice, regardless of treatment onset. Overall, transgenic mice showed a progressive reduction in motor function by P90, with no statistical significance between iFOXO and vehicle treated *hSOD1^G93A^* mice at any time point of treatment (Figure 1b,c,e,f).

**Figure 1.**
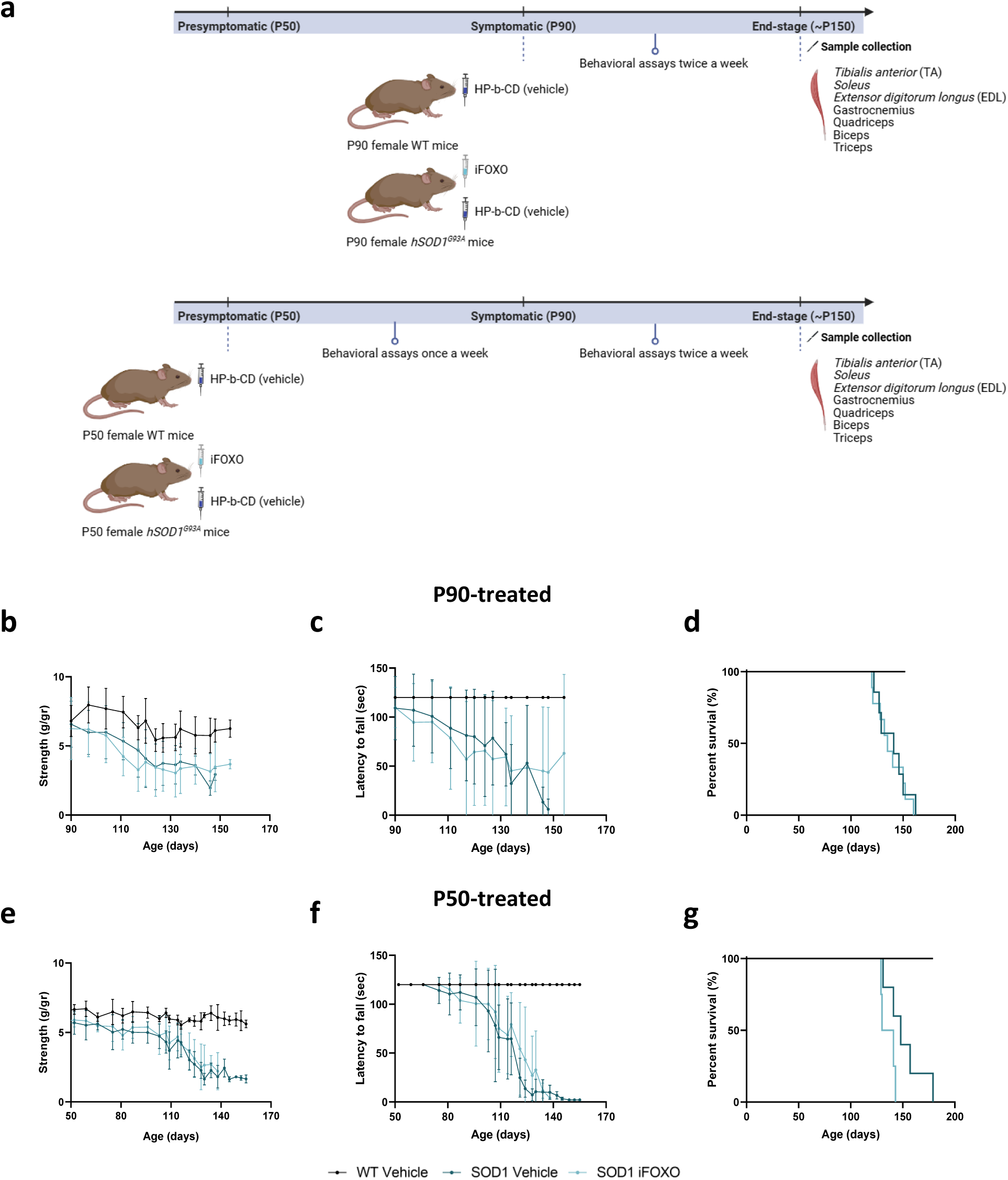
Experimental design and behavioral assays of control, vehicle and iFOXO treated *hSOD1^G93A^*mice starting the treatment at symptomatic and presymptomatic ages. **a)** Experimental timeline of treatment and behavioral assays in *hSOD1^G93A^* mice, including treatment initiation at symptomatic postnatal day 90 (P90) or presymptomatic P50 and sample collection at end-stage. **b)** Grip strength, **c)** wire hanging test and **d)** Kaplan-Meier survival analysis starting the treatment with iFOXO (FOXO inhibitor) at symptomatic P90. **e)** Grip strength, **f)** wire hanging test and **g)** Kaplan-Meier survival analysis starting the treatment with iFOXO at presymptomatic P50. WT, control; SOD1, *hSOD1^G93A^* mice. Data are expressed as mean ± SD. n=5-9/group.

Similarly, iFOXO treatment did not significantly extend the reduced survival of *hSOD1^G93A^* mice. Median survival times were 135 days for iFOXO and 139 days for vehicle treated mice when treatment started at a symptomatic stage (Figure 1d), and 135.5 days for iFOXO and 148 days for vehicle when treatment began at a presymptomatic stage (Figure 1g). These results indicate that daily iFOXO inhibition does not modify overall disease progression in *hSOD1^G93A^*mice.

### Assessment of progressive body mass loss and muscle atrophy in the *hSOD1^G93A^* mice following iFOXO treatment

Longitudinal body mass analysis exhibited a progressive decline of weight in mutant mice, which was significantly lower than age-matched WT controls at end-stage, regardless of treatment onset (Supplementary Figure 1a,b,d,e). Notably, when treatment was initiated at P50, body weight was preserved until P120 in iFOXO treated mice and P135 in vehicle mice, after which a marked decline was observed in both groups (Supplementary Figure 1e).

Weights of various hindlimb and forelimb muscles were measured at end-stage to assess loss of muscle mass. *hSOD1^G93A^* mice displayed significant weight reduction in most analyzed muscles compared to WT controls, regardless of treatment onset. Notably, *soleus* muscles were preserved at both treatment setups, consistent with its well-known atrophy resistance resulting from its high proportion of slow (type I) fibers (Supplementary Figure 1c,f). However, when the treatment began at P50, the *extensor digitorum longus* (EDL) muscle weight was preserved (Supplementary Figure 1f), suggesting a protection of the muscle when treating at the presymptomatic stage. Overall, these findings suggest that iFOXO treatment does not prevent loss of muscle mass in *hSOD1^G93A^* mice, regardless of treatment onset.

### Analysis of muscle histopathological hallmarks in fast- and slow-twitch muscles from *hSOD1^G93A^* mice upon FOXO inhibition

As fast-twitch fibers are more vulnerable in ALS, whereas slow-twitch fibers remain relatively protected (32), fast (TA) and slow (*soleus*) muscles were analyzed to assess histopathological hallmarks associated with ALS progression (Figure 2). In the TA, the average cross-sectional area (CSA) was significantly reduced in *hSOD1^G93A^*mice, indicating marked fiber atrophy. Myofiber size distribution analysis showed a higher proportion of small fibers (500–1,500 μm^2^) in both transgenic groups compared with WT controls, independent of the treatment starting point (Figure 2a,b,d). In contrast, the *soleus* muscle showed no significant differences in mean CSA. However, fiber size distribution exhibited a shift toward smaller fibers in *hSOD1^G93A^* mice (Figure 2a,c,e). The unchanged mean CSA, despite a leftward shift in the distribution, suggests increased fiber-size heterogeneity, with both smaller fibers and preserved large fibers contributing to a stable average. Interestingly, when treatment started at P50, vehicle *hSOD1^G93A^*mice displayed an increased proportion of larger fibers (3,000–4,200 μm^2^), whereas iFOXO treated mice showed a shift toward smaller fibers (Figure 2a,e). These findings confirm that fast-twitch muscles such as TA are more vulnerable to ALS associated degeneration, while slow-twitch muscles like *soleus* are relatively preserved. Nevertheless, iFOXO treatment did not prevent fiber atrophy, regardless of treatment onset.

**Figure 2.**
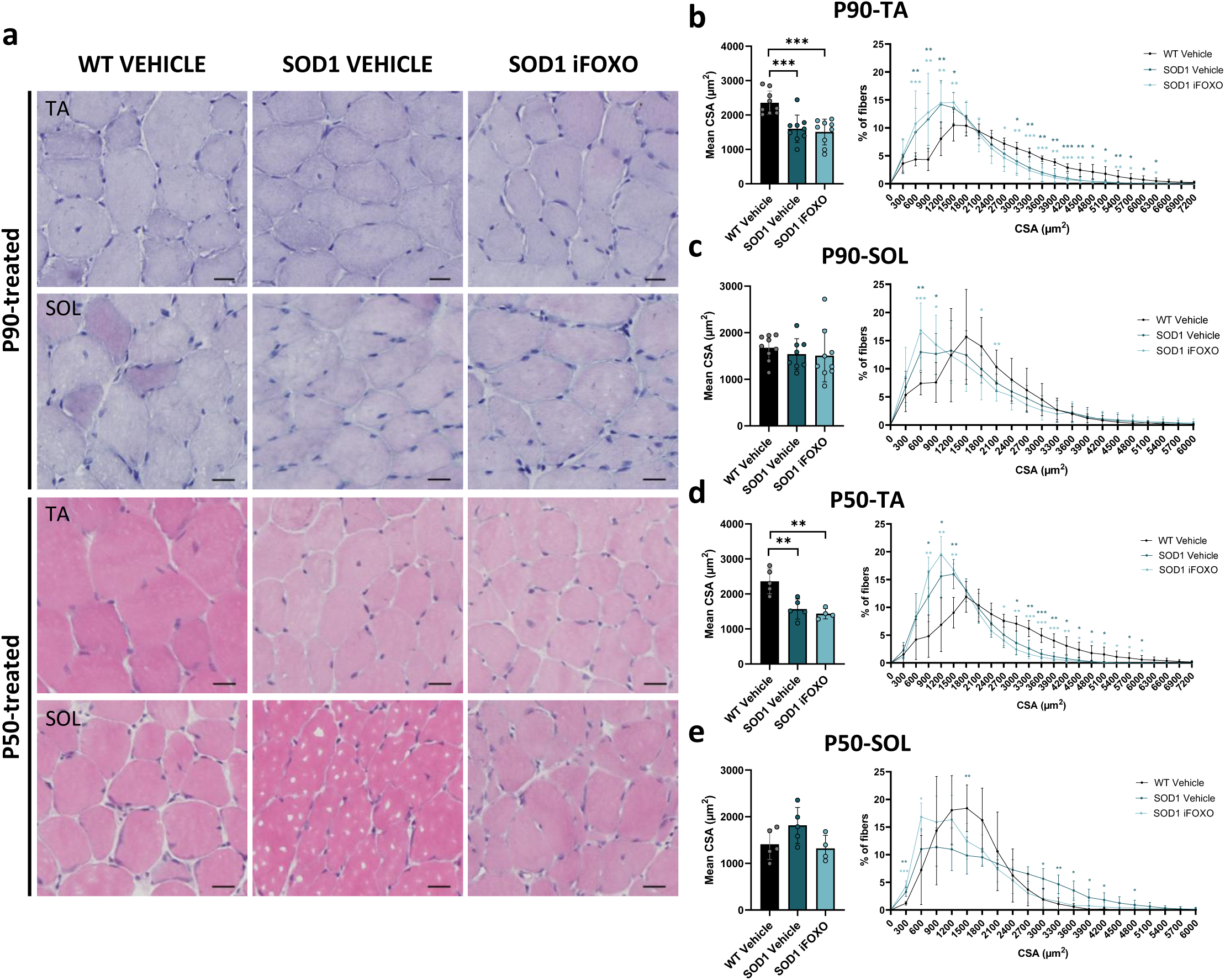
Histopathological characterization of fast and slow muscles of control, vehicle and iFOXO treated *hSOD1^G93A^* mice at end-stage. **a)** Representative images of *hematoxylin and eosin* staining starting the treatment at symptomatic postnatal day 90 (P90) and presymptomatic P50 *hSOD1^G93A^* (SOD1) mice. **b)** Mean fiber cross-sectional area (CSA) and myofiber distribution in the *tibialis anterior* (TA) and **c)** *soleus* (SOL) muscles starting the treatment with iFOXO (FOXO inhibitor) at P90. **d)** Mean fiber CSA and myofiber distribution in the TA and **e)** SOL starting the treatment with iFOXO at P50. WT, control mice. Scale bar: 20 µm. Data are expressed as mean ± SD. n=4-9/group. For normally distributed data with equal variances, one-way ANOVA followed by Tukey’s *post hoc* test was applied. When variances were unequal, Welch ANOVA followed by Games–Howell *post hoc* test was used. For non-normally distributed data, the Kruskal-Wallis test followed by Dunn’s *post hoc* test was applied. Significant differences are indicated as \**p*<0.05, \*\**p*<0.01, \*\*\**p*<0.001.

We next analyzed if this muscle atrophy included some fibrosis or fat infiltration as previously described for ALS disease (42–44). Oil Red O staining revealed no intramuscular fat accumulation, suggesting that muscle degeneration did not progress toward fatty replacement in this model (Supplementary Figure 2). Collagen deposition, assessed by Picrosirius Red staining, was significantly elevated in interstitial and perivascular zones in *hSOD1^G93A^*mice compared with age-matched WT controls in both fast and slow-twitch muscles, consistent with possible pathological extracellular matrix remodeling (Figure 3). iFOXO treatment did not reduce collagen deposition at either treatment onset. Together, these findings confirm that muscle pathology in *hSOD1^G93A^*mice is characterized by fiber atrophy and fibrosis, which iFOXO fails to ameliorate.

**Figure 3.**
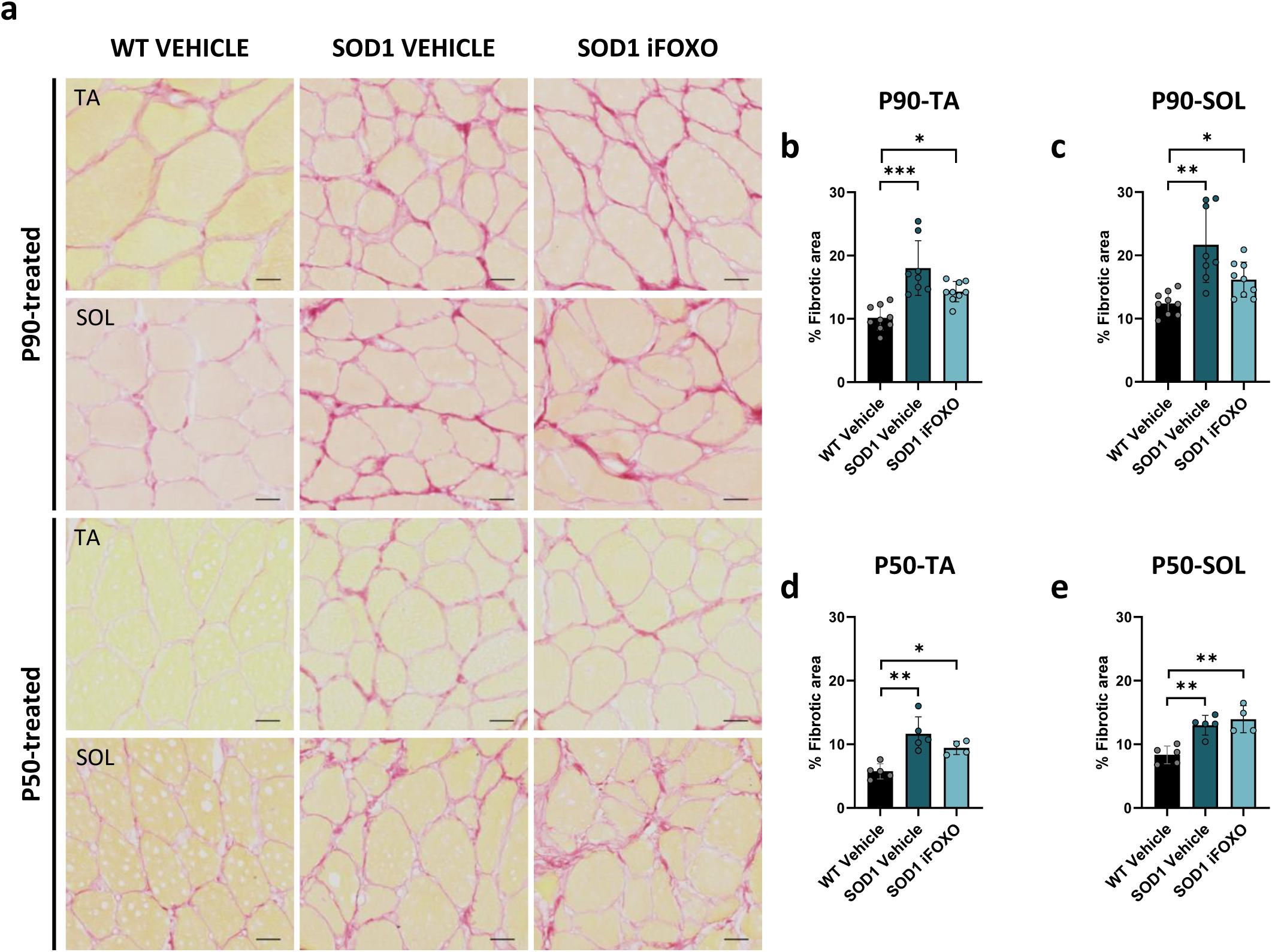
Assessment of fibrosis in fast and slow muscles of control, vehicle and iFOXO treated *hSOD1^G93A^*mice at end-stage. **a)** Representative images of Picrosirius Red staining starting the treatment at symptomatic postnatal day 90 (P90) and presymptomatic P50 *hSOD1^G93A^* (SOD1) mice. **b)** Quantification of the fibrotic area in the *tibialis anterior* (TA) and **c)** *soleus* (SOL) muscles starting the treatment with iFOXO (FOXO inhibitor) at P90. **d)** Quantification of fibrotic area in the TA and **e)** SOL starting the treatment with iFOXO at P90. WT, control mice. Scale bar: 20 µm. Data are expressed as mean ± SD. n=4-9/group. For normally distributed data with equal variances, one-way ANOVA followed by Tukey’s *post hoc* test was applied. When variances were unequal, Welch ANOVA followed by Games–Howell *post hoc* test was used. For non-normally distributed data, the Kruskal-Wallis test followed by Dunn’s *post hoc* test was applied. Significant differences are indicated as \**p*<0.05, \*\**p*<0.01, \*\*\**p*<0.001.

### Impact of iFOXO treatment on muscle stem cells in muscles from *hSOD1^G93A^* mice

Given the presence of muscle atrophy, we examined whether muscle stem cells, the satellite cells (SC), which contributes to muscle maintenance and repair, were affected under these conditions. Dysregulation of SC number or function has been linked in multiple muscle-wasting conditions, including muscular dystrophies and age-related sarcopenia (45). SC number was quantified in TA and *soleus* muscles to assess potential effects of genotype and treatment on the abundance of this population. When treatment was initiated at P90, SC numbers in both TA and *soleus* muscles did not differ between transgenic and WT mice (Figure 4a,b,c), indicating preservation of the satellite cell pool. In contrast, when treatment began at P50, vehicle *hSOD1^G93A^*mice displayed increased SC numbers in the TA compared with WT controls, whereas iFOXO treated mice showed no significant difference (Figure 4a,d). In the *soleus*, *hSOD1^G93A^* mice exhibited elevated SC numbers compared with WT mice, regardless of treatment onset (Figure 4a,e). These results indicate that SC numbers in *hSOD1^G93A^* mice increases in a muscle- and treatment-depending manner, with the most pronounced rise observed in the *soleus*, suggesting a compensatory response aimed at preserving muscle integrity, with no impact of iFOXO treatment on the SC pool.

**Figure 4.**
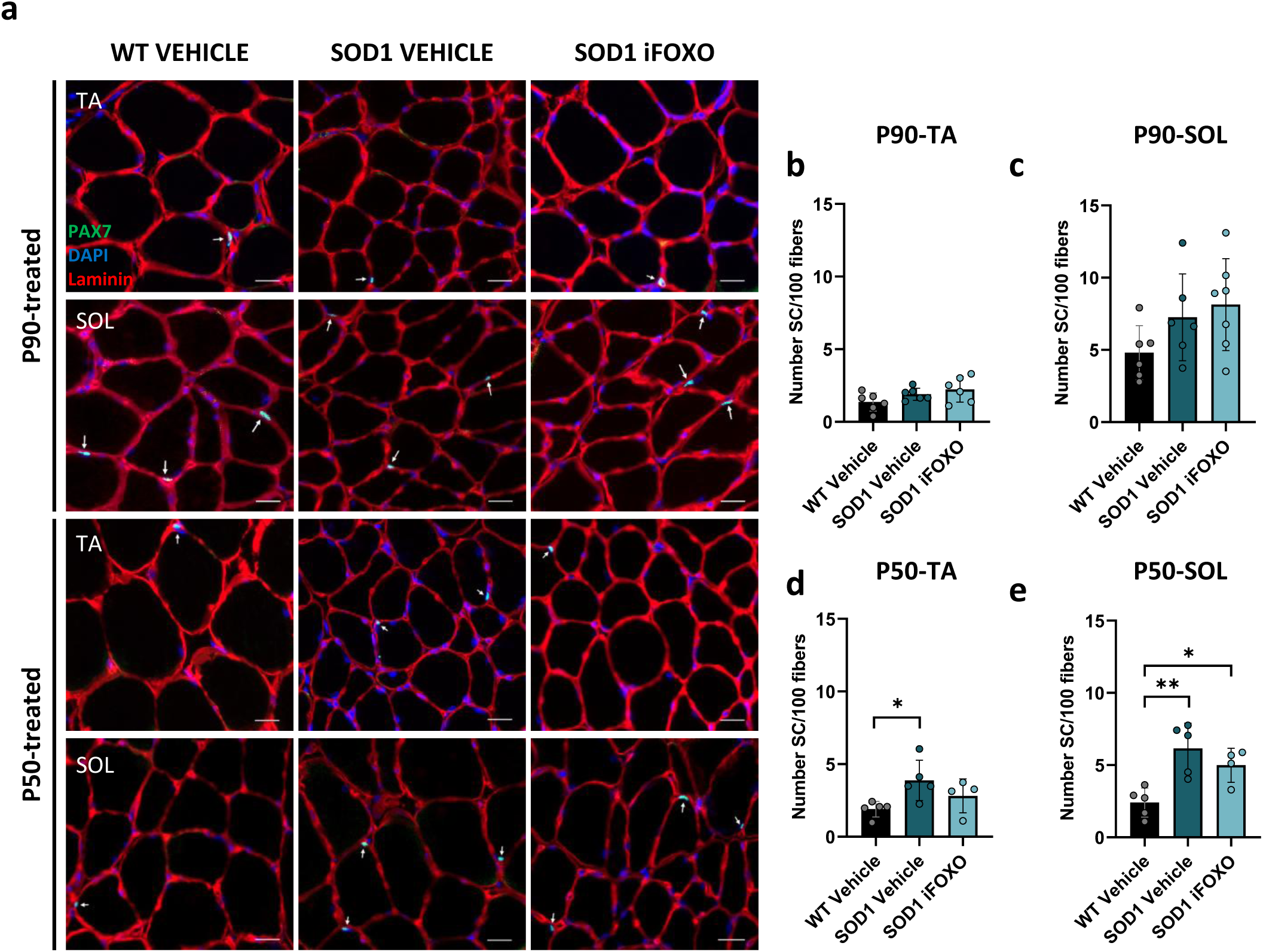
Satellite cells analysis of slow and fast muscles of control, vehicle and iFOXO treated *hSOD1^G93A^*mice at end-stage. **a)** Representative images of PAX7 (green, satellite cell-SC), Laminin (red, basal lamina) and DAPI (blue, nuclei) immunofluorescence. **b)** Number of SC per 100 fibers in the *tibialis anterior* (TA) and **c)** *soleus* (SOL) starting the treatment with iFOXO (FOXO inhibitor) at symptomatic postnatal day 90 (P90). **d)** Number of SC per 100 fibers in the TA and **e)** SOL starting the treatment with iFOXO at presymptomatic P50. WT, control mice. Scale bar: 20 µm. Data are expressed as mean ± SD. n=4-6/group. One-way ANOVA followed by Tukey’s *post hoc* test was applied. Statistical significance is indicated as \**p*<0.05, \*\**p*<0.01, \*\*\**p*<0.001.

### Evaluation of muscle fiber type composition and atrophy after FOXO inhibition in *hSOD1^G93A^* mice

Fiber type composition was also assessed in both fast and slow muscles. Previous studies have already demonstrated that glycolytic fibers are preferentially affected in ALS, whereas oxidative fibers are relatively preserved (32). In the analyzed fast-twitch muscle when treatment started at P90, while fiber proportion was not altered, *hSOD1^G93A^*mice exhibited a strong reduction in type IIB fiber size compared to WT mice, with mild reduction in type IIX and IIA fibers (Figure 5a,b). Strikingly, when treatment started at P50, vehicle *hSOD1^G93A^* mice showed a significant reduction in type IIX fiber proportion with a trend of increasing type IIA, a phenotype recovered with iFOXO treatment (Figure 5a,d). Regarding fiber size, as in the P90 treatment, a strong reduction of type IIB fiber CSA was observed in the P50 treated *hSOD1^G93A^* mice compared to WT mice (Figure 5d). These results indicate that FOXO inhibition does not rescue atrophy across fast fiber subtypes affected in the TA muscle. In the analyzed slow-twitch *soleus* muscle, in mutant animals a metabolic shift toward slow fibers was observed regardless of treatment onset, characterized by an increase in type I fibers and a decrease in type IIA fibers (Figure 5a,c,e). Additionally, *hSOD1^G93A^* mice displayed reduced type IIA fiber size (Figure 5c,e). This shift reflects selective vulnerability of fast fibers, resulting in a relative preservation of slow fibers and more oxidative fiber composition in the *soleus.* Overall, these results indicate that fast-twitch fibers degenerate earlier than slow-twitch fibers in *hSOD1^G93A^* mice, while iFOXO treatment has minimal impact on fiber type composition and fiber-specific atrophy prevention.

**Figure 5.**
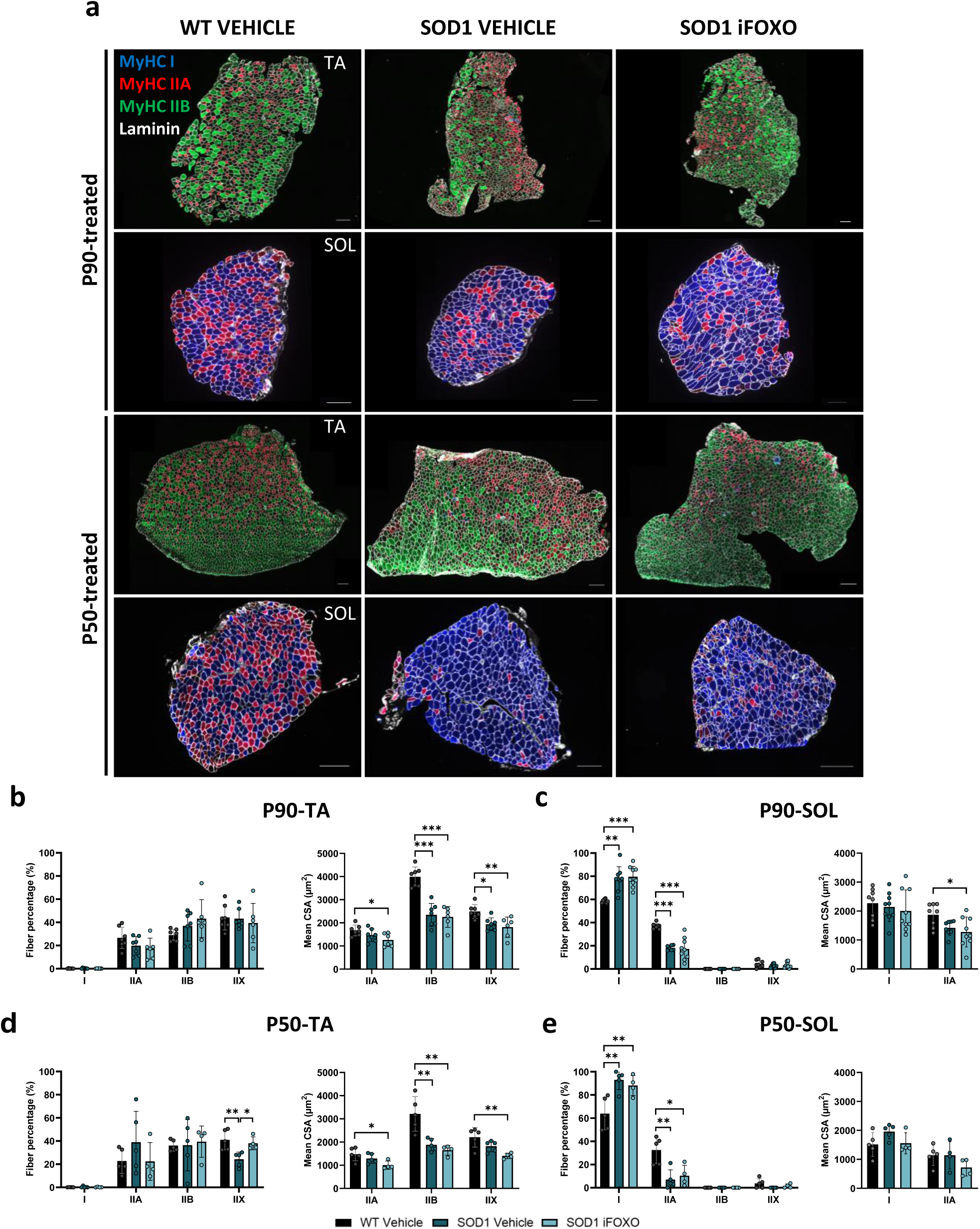
Fiber type composition of fast and slow muscles at end-stage of control, vehicle and iFOXO treated *hSOD1^G93A^* mice at end-stage. **a)** Representative images of myosin heavy chain (MyHC) I (blue), MyHC IIA (red), MyHC IIB (green), MyHC IIX (black), and Laminin (white) immunofluorescence starting the treatment at symptomatic postnatal day 90 (P90) and presymptomatic P50 of control (WT) or *hSOD1^G93A^* (SOD1) mice. Scale bar: 200 µm. **b)** Fiber type proportion and mean fiber cross-sectional area (CSA) in the *tibialis anterior* (TA; fast) and **c)** *soleus* (SOL; slow) starting the treatment with iFOXO (FOXO inhibitor) at P90. **d)** Fiber type proportion and mean fiber CSA in the TA and **e)** SOL starting the treatment with iFOXO at P50. Data are expressed as mean ± SD. n=4-7/group. For normally distributed data with equal variances, one-way ANOVA followed by Tukey’s *post hoc* test was applied. When variances were unequal, Welch ANOVA followed by Games–Howell *post hoc* test was used. For non-normally distributed data, the Kruskal-Wallis test followed by Dunn’s *post hoc* test was applied. Statistical significance is indicated as \**p<*0.05, \*\**p*<0.01, \*\*\**p*<0.001.

### Determination of the effects of iFOXO treatment onset on muscle integrity, adaptive metabolism and stress response pathways

To assess global transcriptional changes in skeletal muscle and determine the contribution of genotype, treatment and treatment starting date, we performed RNA sequencing of quadriceps muscles and posterior principal component analysis (PCA) using the top 500 most variable genes after variance-stabilizing transformation across experimental groups. The first principal component (PC1, 48.2% of variance) segregated samples according to genotype, indicating that genetic differences are the main source of transcriptional variation (Figure 6a). Considering treatment, the PC2 (9.6%), accounted for a smaller portion of variance without showing clear association with any experimental variable, suggesting that iFOXO does not induce strong global changes in gene expression (Figure 6a).

**Figure 6.**
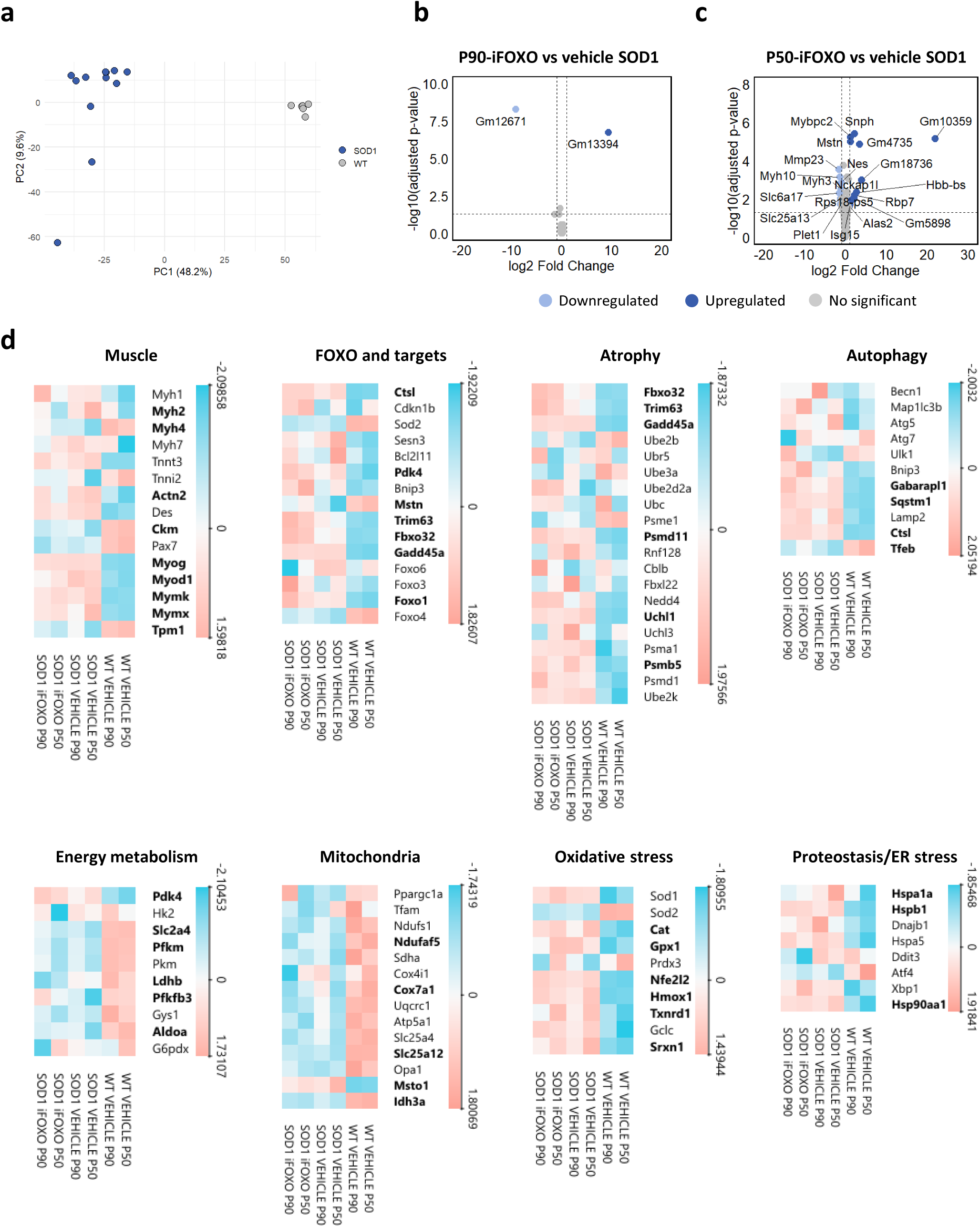
Principal component and differentially expressed genes analysis in quadriceps of control, vehicle and iFOXO treated *hSOD1^G93A^* mice at end-stage. **a)** Principal component analysis (PCA) plot showing genotype as the main source of variance. **b)** Volcano plots comparing DEGs of iFOXO treated SOD1 vs vehicle treated SOD1 mice starting the treatment at P90 and **c)** P50. **d)** Heatmap of the differentially expressed genes (DEG) grouped in eight functional categories: muscle structure, FOXO and targets, atrophy, oxidative stress, autophagy, energy metabolism, mitochondria/oxidative phosphorylation and proteostasis/endoplasmic reticulum (ER) stress. Values in the heatmap are expressed as mean (n=3/group). Genes that are significant in one or more comparison are highlighted in bold. SOD1, *hSOD1^G93A^* mice; P90, starting the treatment at symptomatic postnatal day 90; P50, starting the treatment at presymptomatic postnatal day 50.

Volcano plots, generated to visualize differentially expressed genes (DEG), revealed 738 upregulated (UR) and 544 downregulated (DR) genes when treatment started at P90 in vehicle *hSOD1^G93A^* mice compared to WT, while iFOXO treatment induced 947 UR and 687 DR genes (Supplementary Figure 3a,b). When treatment began at P50, vehicle mice showed 1,094 UR and 842 DR genes, whereas iFOXO treated mice 775 UR and 566 DR genes compared to WT mice (Supplementary Figure 3c,d). As iFOXO reduced the number of DEGs when initiated at P50, but increased them when started at P90, direct comparison between iFOXO treated mice at these two treatment-starting points showed no significant DEGs (Supplementary Figure 3e). While differential expression analysis between iFOXO and vehicle mice reported only two pseudogenes when treatment began at P90 (Figure 6b), treatment initiated at P50 presented 12 UR (5 pseudogenes) and 8 DR genes. DR genes included *Mmp23*, *Myh10*, *Myh3*, *Slc6a17*, *Slc25a13*, *Plet1*, *Nckap1l* and *Nes*, many of which are associated with muscle structure, metabolism and progenitor cell function. UR genes include *Mstn*, *Mybpc2*, *Isg15*, *Snph*, *Hbb-bs*, *Rbp7* and *Alas2*, suggesting a subtle modulation on sarcomeric proteins, stress response and metabolic pathways (Figure 6c).

Functional enrichment analysis confirmed genotype as the primary driver of transcriptional changes in the skeletal muscle (Supplementary Figure 4). Indeed, DR pathways in SOD1-G93A muscles when treatment started at the symptomatic age, common for vehicle or iFOXO (Supplementary Figure 4a,b), showed global reduction in muscle activity, metabolic energy production, and adaptive signaling capacity. These data indicate signs of weakened muscle function, diminished canonical growth signaling, suppressed metabolic flux, and diminished ability to sustain energy demanding activity. Conversely, UR genes were enriched in pathways related to contractile and adaptive demands, compensatory regeneration, apoptosis and oxidative stress metabolism (Supplementary Figure 4c,d). This UR signaling indicates a state of hyperactive muscle remodeling, with the skeletal muscle under high stress. Together, SOD1 mutant muscles appeared simultaneously overactivated in structural turnover yet energetically compromised, creating a state of compensatory regeneration under metabolic constraint. Considering treatment starting at the presymptomatic stage, DR signaling pathways showed again broad reduction in energy generation, metabolic processing, and contractile function, characterized by impaired glucose usage, weakened oxidative metabolism, reduced calcium dynamics, and compromised muscle performance (Supplementary Figure 4e,f). However, UR pathways point to a hyper regenerative, stress responsive phenotype, where the muscle attempts to maintain function despite increased metabolic and oxidative demands (Supplementary Figure 4g,h). Collectively and independent of treatment onset, SOD1 mutants present a muscle environment under chronic stress that attempts to preserve its function via heightened apoptotic turnover and transcriptional activation of myogenic fusion/differentiation and oxidative stress programs. Yet, it remains constrained by deficient contractile support and a hypometabolic energy landscape.

We next analyzed treatment effect, confirming that iFOXO treatment modulated several pathways in a treatment onset dependent manner. When treatment was initiated at P90, iFOXO treatment DEG pathways define a muscle environment in which mitochondrial oxidative energy production and Ca²⁺ homeostasis are impaired, while stress responsive, insulin/ketone sensing, and proliferative signaling programs become activated (Supplementary Figure 4b,d). This pattern suggests that iFOXO induces a compensatory attempt to maintain cellular function under metabolic limitation, promoting oxidative stress responses and cytoskeletal remodeling rising in parallel to energy production deficits. The muscle appears to activate signaling pathways associated with adaptation, yet on a foundation of reduced mitochondrial output and impaired calcium regulation. When treatment began at P50, iFOXO push the muscle to adapt via mitochondrial repositioning, alternative fuel uptake, and oxidative stress responses, yet does so while core structural assembly, reserve energy metabolism, and key remodeling signals are suppressed (Supplementary Figure 4f,h). Again, this suggests that iFOXO promotes compensatory metabolic/organellar adaptation on a weakened structural and signaling foundation, predisposing to incomplete or non-productive functional compensation under physiological stress. These results indicate that while genotype is the main determinant of transcriptional changes, iFOXO produces a treatment onset dependent modulation. In summary, treatment at P90 enhanced a compensatory but incomplete adaptative response aiming at preserving muscle function despite underlying metabolic and calcium-handling deficits. Yet, treatment at P50 tried to compensate functional adaptation but on a weakened structural and signaling foundation.

### Differential gene expression in mutant mice following iFOXO treatment

Considering the top modified signaling pathways, we defined eight functional categories: muscle structure, FOXO and targets, atrophy, oxidative stress, autophagy, energy metabolism, mitochondria/oxidative phosphorylation and proteostasis/endoplasmic reticulum (ER) stress. Muscle-related transcripts were mainly UR in mutant mice, including key regulators of myogenesis (*Myog*, *Myod1, Mymk* and *Mymx*). Notably, genes encoding contractile components showed differential patterns: fast oxidative myosin *Myh2* was further UR only in vehicle samples, while fast glycolytic *Myh4* was DR in mutant mice, with the exception of iFOXO treated mice when treatment initiated at P90. These findings are consistent with activation of myogenic programs and ongoing muscle remodeling, indicating that iFOXO at the symptomatic stage may contribute to the relative preservation of vulnerable fast glycolytic fibers. Interestingly, *Foxo1*, together with several canonical downstream targets (*Pdk4*, *Fbxo32* (atrogin-1), *Trim63* (MuRF1), *Ctsl, Gadd45a*), were also UR in iFOXO treated mice. This likely reflects a compensatory response to FOXO inhibition rather than direct activation of FOXO pathway itself (Figure 6d).

Atrophy related genes, such as atrogin-1 and MuRF1 and components of the ubiquitin/proteasome system were UR, indicating that FOXO inhibition alone may be not sufficient to fully suppress muscle wasting pathways. Autophagy related genes, including *Gabarapl1*, *Ctsl* and *Sqstm1*, were elevated, suggesting activation of multiple steps of the autophagic pathway and potential changes in autophagic flux (Figure 6d).

Genes involved in energy metabolism and mitochondrial function appeared predominantly DR, consistent with the metabolic impairment of *hSOD1^G93A^*mice: (i) oxidative phosphorylation (*Idh3a*, *Cox7ai* and *Ndufaf5*) DR indicates compromised mitochondrial activity and energy production; (ii) *Pdk4* UR reflects a metabolic shift promoting fatty acid oxidation over glucose utilization; (iii) glycolytic and glucose transport (*Pfkm*, *Ldhb, Pfkfb3, Aldoa*, *Slc2a4/Glut4*) DR suggests impaired glucose utilization. This profile supports a shift toward a more oxidative and energetically inefficient state, reflecting impaired mitochondrial function and reduced glucose use, a hallmark of ALS-associated muscle pathology. However, FOXO inhibition fails to rescue the persistent energetic deficit observed in ALS muscle (Figure 6d).

Oxidative stress related genes were predominantly UR (*Cat*, *Gpx1*, *Nfe2l2/Nrf2*, *Hmox1*) indicating enhanced antioxidant response. Markers of proteostasis/ER stress (*Hspb1*, *Hsp90aa1*) were UR, whereas *Atf4* and *Ddit3* (*Chop*) levels did not increase, reflecting a moderate and adaptive activation of cellular stress response (Figure 6d).

Overall, iFOXO and vehicle *hSOD1^G93A^* mice present similar transcriptional profiles: UR of genes related to muscle structure, FOXO targets, atrophy, autophagy, oxidative stress and proteostasis/ER stress, and DR of those involved in energy metabolism, glycolysis, mitochondrial function and oxidative phosphorylation, consistent with genotype differences.

## DISCUSSION

Based on its previously established ability to restore myogenic defects *in vitro* and to improve locomotor and survival outcomes in *Drosophila* ALS models (19), we assessed daily iFOXO treatment and found that iFOXO alone does not rescue the disease phenotype in *hSOD1^G93A^*female mice. Though this treatment has previously been shown to be effective at preventing muscle fiber atrophy in cachexia and diabetic models (31,46,47), our data indicate that targeting FOXO systemically is not sufficient to stop disease progression in this model. Importantly, iFOXO did not exacerbate pathology, suggesting a safe pharmacological profile if considered in combination with other strategies.

Assessment of body mass and muscle weights revealed progressive weight loss and significant atrophy in multiple hindlimb and forelimb muscles in *hSOD1^G93A^* mice. However, the relative preservation of *soleus* muscles is consistent with its predominantly oxidative fiber composition and its reported resistance to ALS degeneration (32). Noticeably, preservation of EDL muscle weight when treatment was initiated at P50, suggests that early interventions can induce long-lasting protection to fast glycolytic muscles, as once muscle atrophy is established (P90) cannot be restored. Histopathological analysis further corroborates these observations, showing pronounced fiber atrophy in fast-twitch muscles and increased fibrosis in both fast- and slow-twitch muscles. This increase in collagen deposition reflects pathological extracellular matrix remodeling, a feature also reported in ALS (48,49). Finally, stem SC population increases at earlier intervention, consistent with an ineffective compensatory response to muscle stress. As previously observed (32), fiber type analyses revealed preferential degeneration of fast-twitch fibers, reducing type IIB/IIX fibers, whereas slow-twitch fibers were preserved.

Transcriptomic profiling analyses reveal that genotype is the dominant driver of gene expression changes in *hSOD1^G93A^* mice skeletal muscle, with iFOXO treatment producing only subtle, treatment onset dependent effects. This observation aligns with previous studies showing that ALS itself profoundly reshapes muscle transcriptional programs (50). Despite the absence of significant DEGs between iFOXO P90 vs. P50-treated *hSOD1^G93A^* mice, the transcriptional profiles do diverge from WT mice depending on treatment onset. Treatment at P50 resulted in a lower DEGs compared to P90, suggesting that early FOXO inhibition does not establish an alternative transcriptional profile but instead dampens the extent of transcriptional remodeling associated with disease progression.

Consistent with a chronic stress state, functional enrichment analysis in *hSOD1^G93A^* mice exhibits a transcriptional landscape characterized by impaired energy metabolism, contractile function, and adaptive signaling, together with activation of pathways related to stress responses, apoptosis and muscle remodeling. Though transcriptional programs linked to muscle repair were activated, the absence of histological evidence of effective regeneration suggests ineffective repair response. Indeed, persistent oxidative stress and activation of apoptotic pathways may impair regeneration, ultimately contributing to muscle atrophy. iFOXO treatment does not reverse the underlying transcriptional alterations but modulates metabolic and structural pathways in a treatment onset dependent manner. Supporting metabolic adaptation in an advanced disease context, P90 enhanced metabolic flexibility, including insulin and ketone responses. When initiated at P50, iFOXO promoted lipid utilization, ketone metabolism and mitochondrial organization, suggesting a shift toward a more oxidative metabolic profile under sustained muscle stress, consistent with the observed histological findings.

Though subtle, differential expression and functional enrichment analyses indicate that iFOXO treatment can modulate specific muscle-related pathways. Although myogenic regulators were UR, contractile genes displayed stage dependent responses. Fast glycolytic myosin *Myh4*, more vulnerable in ALS, was DR in most conditions but preserved after iFOXO treatment when initiated at P90, suggesting a potential protective effect. Nevertheless, pathways involved in atrophy, autophagy, proteostasis/ER stress remained largely UR in both iFOXO and vehicle treated *hSOD1^G93A^* mice, indicating that FOXO inhibition alone is not sufficient to fully reverse these molecular alterations.

Our results suggest that the timing of iFOXO intervention influences muscle responses in *hSOD1^G93A^* mice. This highlights the importance of considering treatment onset when planning therapeutic strategies, as there may be an optimal window during which the muscle plasticity allows the treatment to be most effective.

An unexpected observation was the biological activity of the vehicle. HP-b-CD is commonly used as an adjuvant for therapeutic agents to aid solubility and it has been described to not alter disease progression in *hSOD1^G93A^*mice, even at big doses (51). However, HP-b-CD preserved body weight, extended life expectancy and produced a shift toward larger fibers in the *soleus* of *hSOD1^G93A^* mice when treatment was initiated at P50. Cyclodextrins are known to modulate cholesterol availability (52,53), which may alter lipid metabolism and membrane associated signaling. Despite these effects, the *soleus* retained its characteristic oxidative profile, supporting the preservation of overall physiological muscle traits.

It is worth noting that this study was conducted in the *hSOD1^G93A^*mouse model, a well characterized and widely used ALS model. We have previously observed beneficial effects of iFOXO in TDP-43 and FUS *Drosophila* models, as well as patient-derived primary muscle cells (19). *hSOD1^G93A^* mouse model was chosen due to its robustness and reproducibility, whereas TDP-43 and FUS mouse models present limitations and do not recapitulate disease symptoms. Therefore, the limited impact of iFOXO in *hSOD1^G93A^* mice may reflect model specific differences in disease mechanisms or in the responsiveness of skeletal muscle to FOXO inhibition, suggesting that testing this treatment in additional ALS models could reveal effects not captured in the SOD1 context.

## CONCLUSIONS

Our results indicate that iFOXO exerts subtle, stage-depending effects on skeletal muscle transcriptional programs, slightly enhancing metabolic adaptive and oxidative stress responses; but does not prevent motor deficits, weight loss, muscle atrophy or fiber specific degeneration in *hSOD1^G93A^*female mice. As FOXO inhibition alone is not sufficient to overcome the genotype driven alterations in ALS skeletal muscle, combinatorial approaches targeting multiple pathways may be required to achieve meaningful therapeutic benefits. These findings provide a mechanistic framework for future studies exploring FOXO based interventions in neuromuscular disorders characterized by degeneration, metabolic imbalance and oxidative stress, and highlight the importance of considering both treatment onset and vehicle effects in preclinical therapeutic studies.

### Limitations of the study

This study relied on a single genetic model, the SOD1-G93A mutation, which although representing a mutation found in some sporadic ALS cases, may not reflect the full genetic and phenotypic diversity of the disease. Previous work demonstrated beneficial effects of iFOXO in muscle-specific knockdown models of TDP-43 and FUS in *Drosophila*, suggesting that results obtained from the *hSOD1^G93A^* mouse model may not fully translate to other ALS contexts. The exclusive use of female mice limits the applicability of our conclusions, as sex-related differences are known to influence disease onset, progression, and response to interventions. Future studies should consider both sexes to better understand potential variability in treatment outcomes. Finally, a notable limitation is that the vehicle, HP-b-CD, itself exerts effects on body weight, survival, and muscle fiber composition in our study. While previous reports indicated that HP-b-CD may not significantly alter disease progression in this model, our observations suggest that vehicle related effects must be considered in preclinical therapeutic studies.

## Authors’ contribution

This project was administered by SA-M. SA-M conceived, planned and supervised these experiments. AV-G, IA, ML, AE, OP-M, MR-C, AO, LM-M, CR-R, AV, BD and MG-P conducted laboratory experiments. AV-G, IA and MR-H performed data analysis. LM-M and RO provided mice for the experiments. IV, AM, AS, ALM and SA-M contributed to funding acquisition and supported the project. AV-G and SA-M wrote, reviewed and edited the original draft. All authors have read, reviewed and approved the final version of this paper.

