## Supplementary Figure 1 for "Comprehensive characterization of skeletal muscle remodeling in *hSOD1^G93A^* mice reveals limited functional impact of systemic FOXO1 inhibition"

### Slide 1
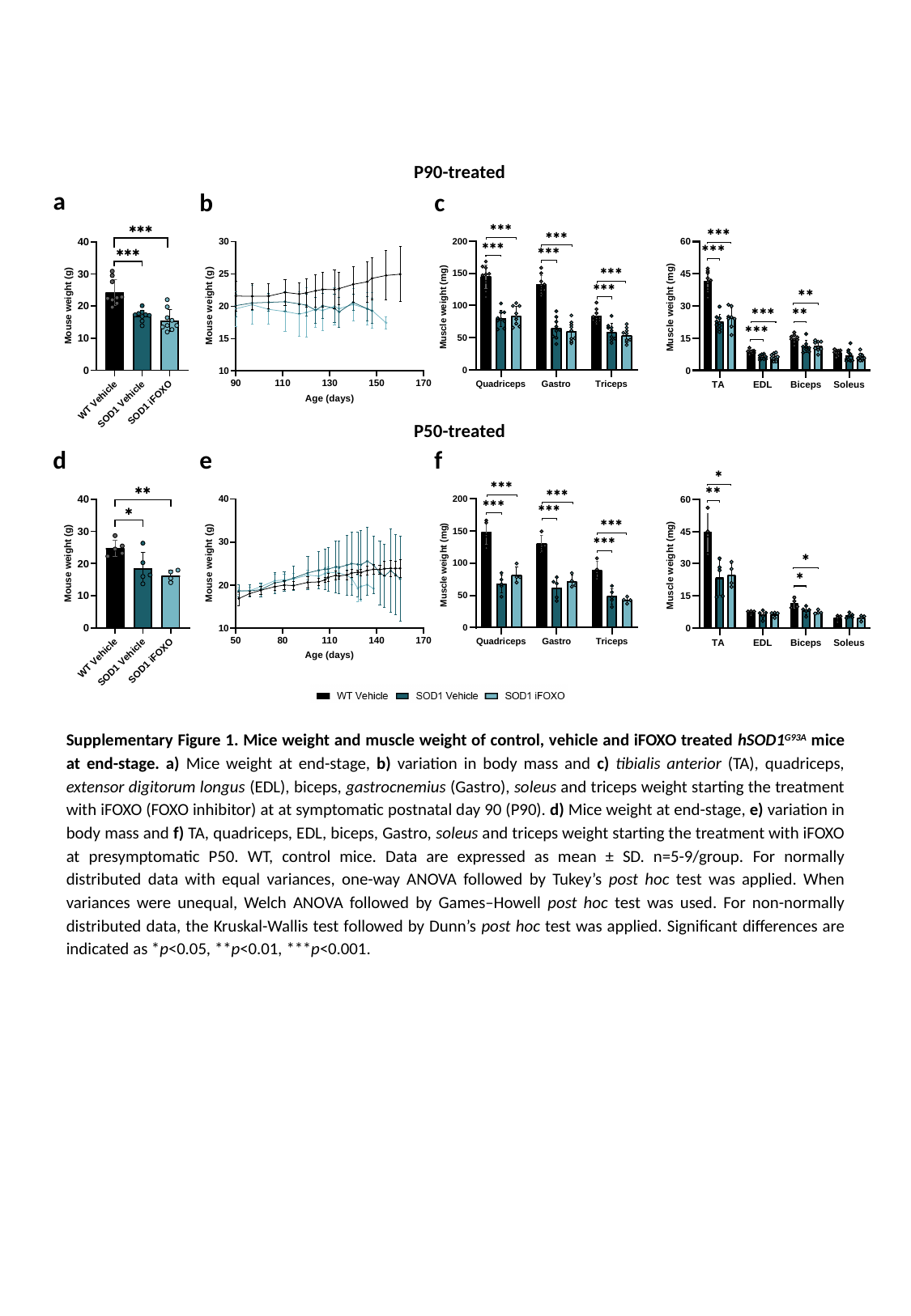

P90-treated
a
b
c
P50-treated
d
e
f
Supplementary Figure 1. Mice weight and muscle weight of control, vehicle and iFOXO treated hSOD1^G93A^ mice at end-stage. a) Mice weight at end-stage, b) variation in body mass and c) tibialis anterior (TA), quadriceps, extensor digitorum longus (EDL), biceps, gastrocnemius (Gastro), soleus and triceps weight starting the treatment with iFOXO (FOXO inhibitor) at at symptomatic postnatal day 90 (P90). d) Mice weight at end-stage, e) variation in body mass and f) TA, quadriceps, EDL, biceps, Gastro, soleus and triceps weight starting the treatment with iFOXO at presymptomatic P50. WT, control mice. Data are expressed as mean ± SD. n=5-9/group. For normally distributed data with equal variances, one-way ANOVA followed by Tukey’s post hoc test was applied. When variances were unequal, Welch ANOVA followed by Games–Howell post hoc test was used. For non-normally distributed data, the Kruskal-Wallis test followed by Dunn’s post hoc test was applied. Significant differences are indicated as *p<0.05, **p<0.01, ***p<0.001.
