## Supplementary Figure 2 for "Comprehensive characterization of skeletal muscle remodeling in *hSOD1^G93A^* mice reveals limited functional impact of systemic FOXO1 inhibition"

### Slide 1
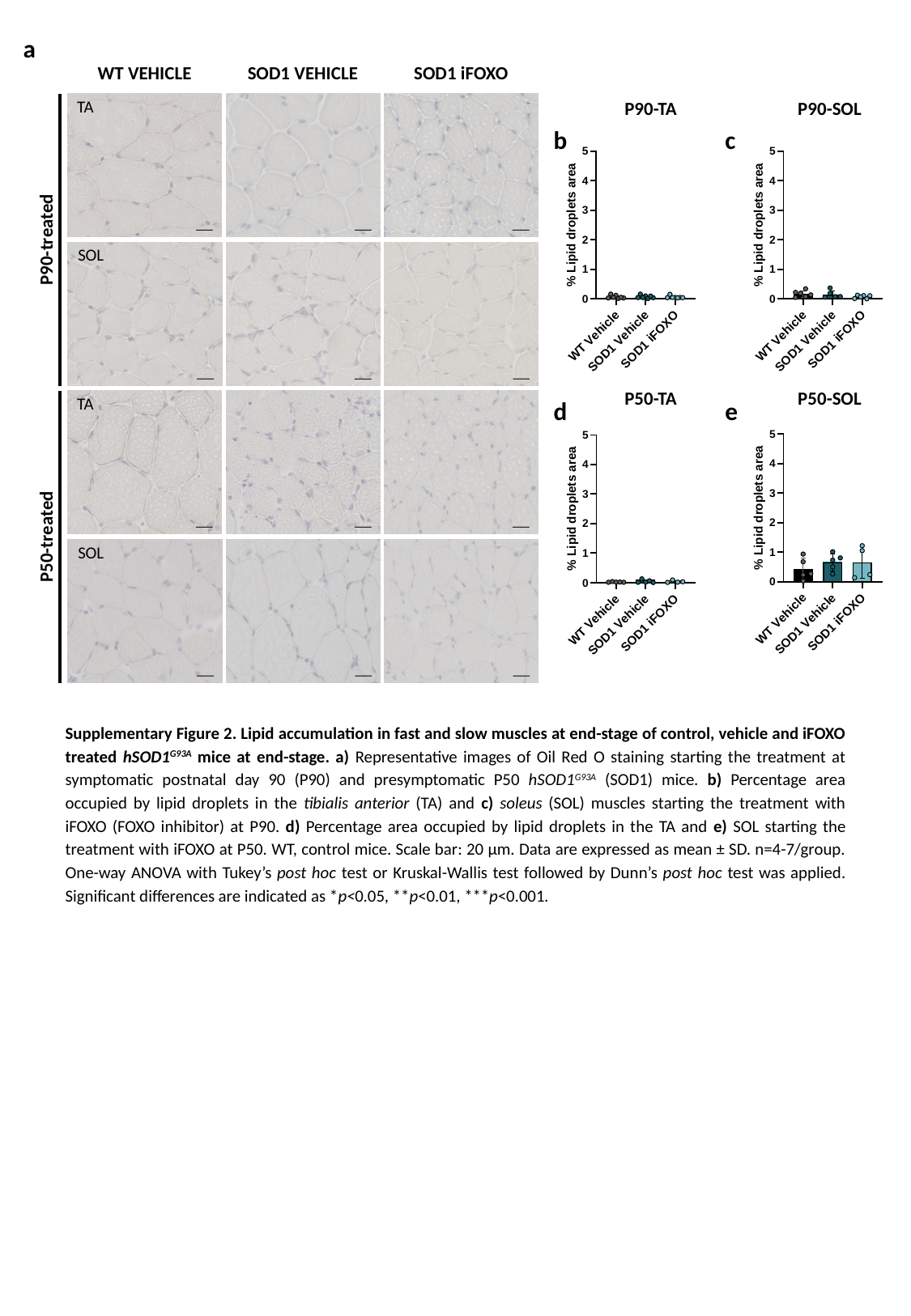

a
WT VEHICLE
SOD1 VEHICLE
SOD1 iFOXO
TA
P90-TA
P90-SOL
b
c
P90-treated
SOL
P50-TA
P50-SOL
d
e
TA
P50-treated
SOL
Supplementary Figure 2. Lipid accumulation in fast and slow muscles at end-stage of control, vehicle and iFOXO treated hSOD1^G93A^ mice at end-stage. a) Representative images of Oil Red O staining starting the treatment at symptomatic postnatal day 90 (P90) and presymptomatic P50 hSOD1^G93A^ (SOD1) mice. b) Percentage area occupied by lipid droplets in the tibialis anterior (TA) and c) soleus (SOL) muscles starting the treatment with iFOXO (FOXO inhibitor) at P90. d) Percentage area occupied by lipid droplets in the TA and e) SOL starting the treatment with iFOXO at P50. WT, control mice. Scale bar: 20 µm. Data are expressed as mean ± SD. n=4-7/group. One-way ANOVA with Tukey’s post hoc test or Kruskal-Wallis test followed by Dunn’s post hoc test was applied. Significant differences are indicated as *p<0.05, **p<0.01, ***p<0.001.
