## Supplementary Figure 3 for "Comprehensive characterization of skeletal muscle remodeling in *hSOD1^G93A^* mice reveals limited functional impact of systemic FOXO1 inhibition"

### Slide 1
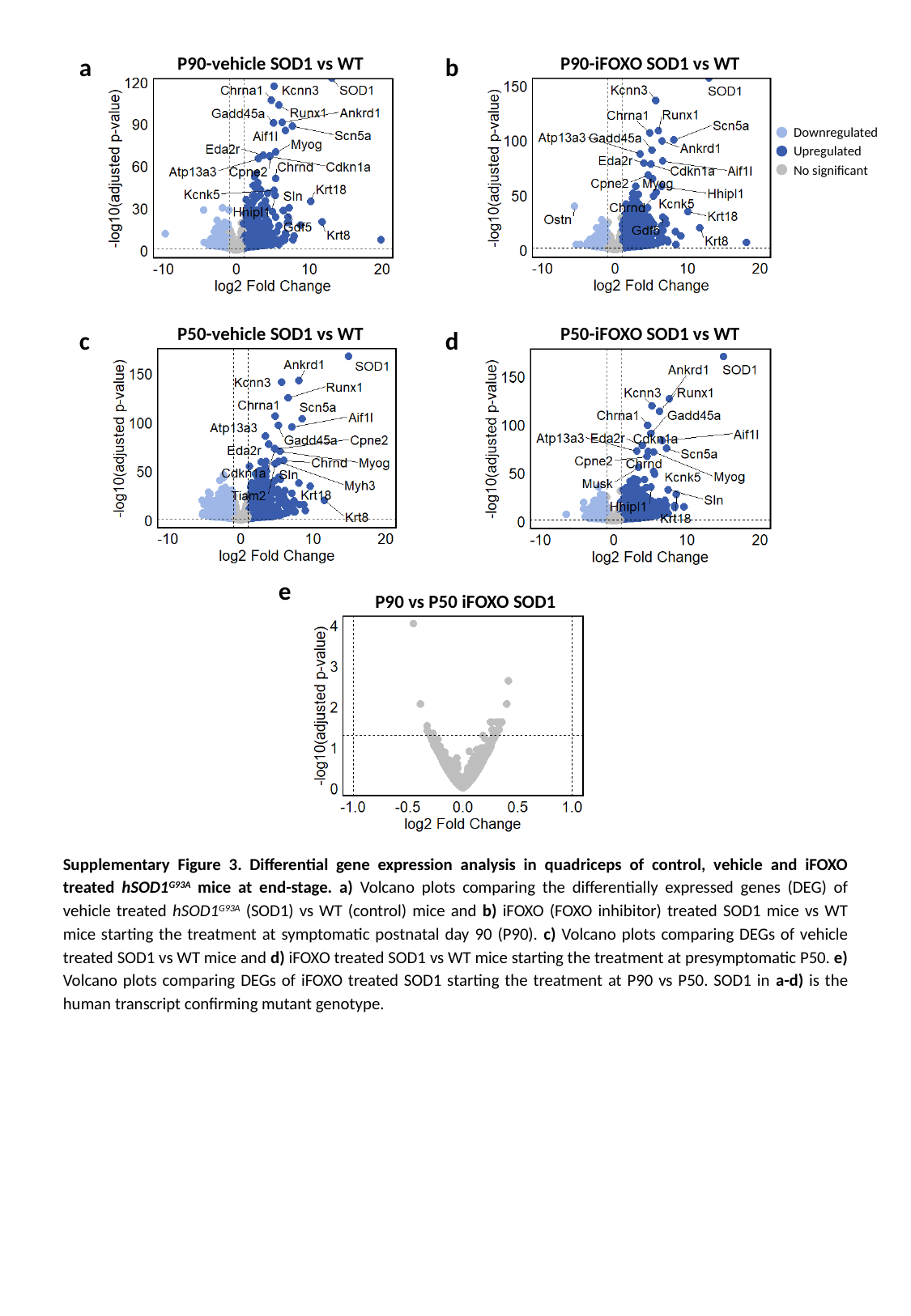

a
b
P90-vehicle SOD1 vs WT
P90-iFOXO SOD1 vs WT
Downregulated
Upregulated
No significant
c
d
P50-vehicle SOD1 vs WT
P50-iFOXO SOD1 vs WT
e
P90 vs P50 iFOXO SOD1
Supplementary Figure 3. Differential gene expression analysis in quadriceps of control, vehicle and iFOXO treated hSOD1^G93A^ mice at end-stage. a) Volcano plots comparing the differentially expressed genes (DEG) of vehicle treated hSOD1^G93A^ (SOD1) vs WT (control) mice and b) iFOXO (FOXO inhibitor) treated SOD1 mice vs WT mice starting the treatment at symptomatic postnatal day 90 (P90). c) Volcano plots comparing DEGs of vehicle treated SOD1 vs WT mice and d) iFOXO treated SOD1 vs WT mice starting the treatment at presymptomatic P50. e) Volcano plots comparing DEGs of iFOXO treated SOD1 starting the treatment at P90 vs P50. SOD1 in a-d) is the human transcript confirming mutant genotype.
