## Supplementary Figure 4 for "Comprehensive characterization of skeletal muscle remodeling in *hSOD1^G93A^* mice reveals limited functional impact of systemic FOXO1 inhibition"

### Slide 1
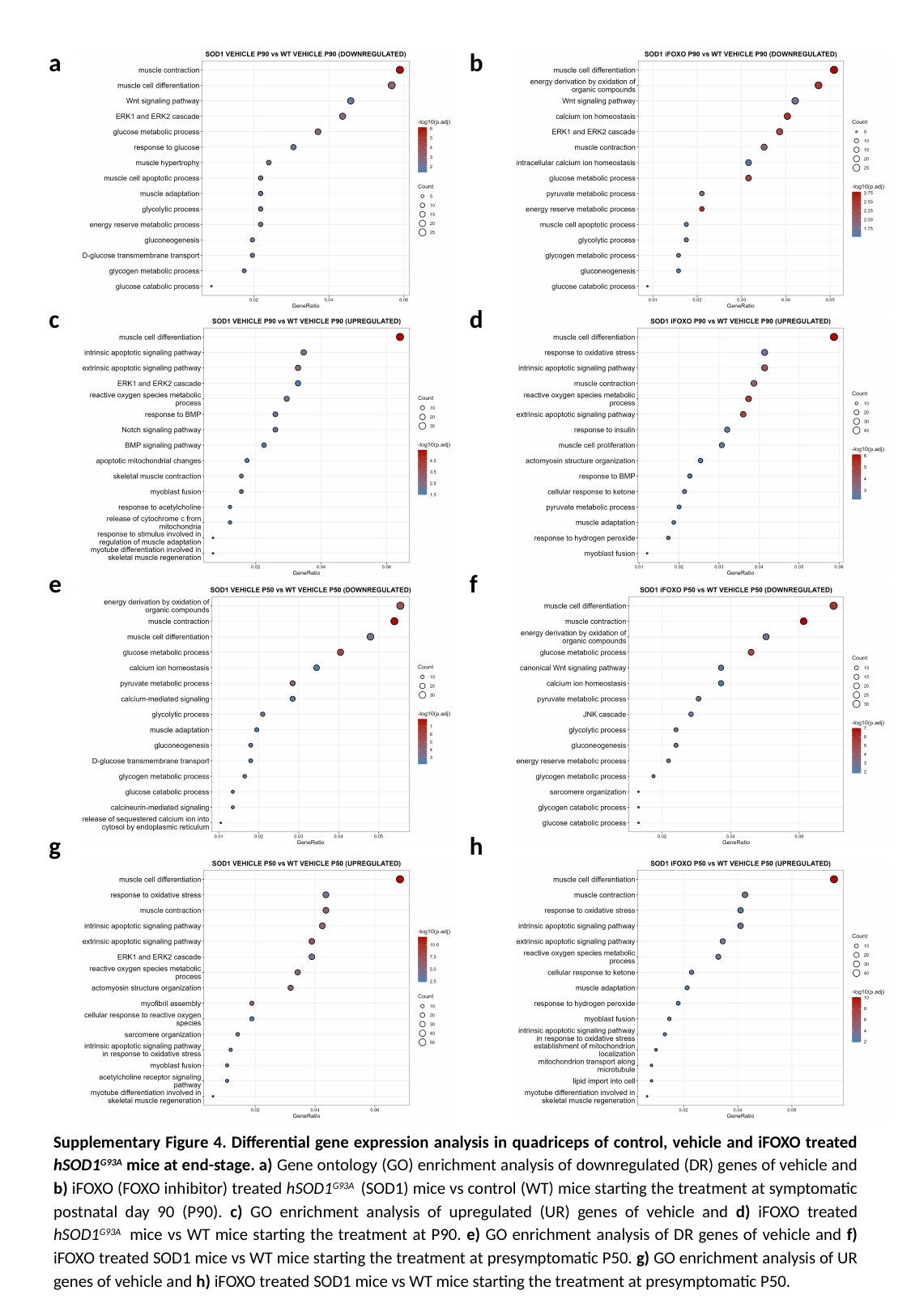

a
b
c
d
e
f
g
h
Supplementary Figure 4. Differential gene expression analysis in quadriceps of control, vehicle and iFOXO treated hSOD1^G93A^ mice at end-stage. a) Gene ontology (GO) enrichment analysis of downregulated (DR) genes of vehicle and b) iFOXO (FOXO inhibitor) treated hSOD1^G93A^ (SOD1) mice vs control (WT) mice starting the treatment at symptomatic postnatal day 90 (P90). c) GO enrichment analysis of upregulated (UR) genes of vehicle and d) iFOXO treated hSOD1^G93A^ mice vs WT mice starting the treatment at P90. e) GO enrichment analysis of DR genes of vehicle and f) iFOXO treated SOD1 mice vs WT mice starting the treatment at presymptomatic P50. g) GO enrichment analysis of UR genes of vehicle and h) iFOXO treated SOD1 mice vs WT mice starting the treatment at presymptomatic P50.
